# Differential transcript usage from RNA-seq data: isoform pre-filtering improves performance of count-based methods

**DOI:** 10.1101/025387

**Authors:** Charlotte Soneson, Katarina L. Matthes, Malgorzata Nowicka, Charity W. Law, Mark D. Robinson

**Affiliations:** Institute of Molecular Life Sciences, University of Zurich, Switzerland; SIB Swiss Institute of Bioinformatics, Switzerland; Division of Chronic Disease Epidemiology, Epidemiology, Biostatistics and Prevention Institute (EPBI), University of Zurich, Switzerland; Cancer Registry Zurich and Zug, University Hospital Zurich, Switzerland; Molecular Medicine Division, Walter and Eliza Hall Institute of Medical Research, Australia

**Author notes:** Equal contribution (Charlotte Soneson, Katarina L. Matthes).

## Abstract

Large-scale sequencing of cDNA (RNA-seq) has been a boon to the quantitative analysis of transcriptomes. A notable application is the detection of changes in transcript usage between experimental conditions. For example, discovery of pathological alternative splicing may allow the development of new treatments or better management of patients. From an analysis perspective, there are several ways to approach RNA-seq data to unravel differential transcript usage, such as annotation-based exon-level counting, differential analysis of the ‘percent spliced in’ measure or quantitative analysis of assembled transcripts. The goal of this research is to compare and contrast current state-of-the-art methods, as well as to suggest improvements to commonly used workflows.

We assess the performance of representative workflows using synthetic data and explore the effect of using non-standard counting bin definitions as input to a state-of-the-art inference engine (DEXSeq). Although the canonical counting provided the best results overall, several non-canonical approaches were as good or better in specific aspects and most counting approaches outperformed the evaluated event- and assembly-based methods. We show that an incomplete annotation catalog can have a detrimental effect on the ability to detect differential transcript usage in transcriptomes with few isoforms per gene and that isoform-level pre-filtering can considerably improve false discovery rate (FDR) control.

Count-based methods generally perform well in detection of differential transcript usage. Controlling the FDR at the imposed threshold is difficult, mainly in complex organisms, but can be improved by pre-filtering of the annotation catalog.

## Background

High-throughput sequencing of cDNA fragment populations, commonly known as RNA-seq, has rapidly become a default standard for profiling the composition of the transcriptome and quantify the expression of individual transcriptional units (such as genes, transcripts or exons). One of the key analysis challenges with RNA-seq data is to infer a set of such units that change their expression level or expression pattern between conditions. For example, identifying transcriptional changes that occur between normal and disease states may lead to markers of progression or prognosis, knowledge of molecules that can be pharmaceutically corrected, or to an understanding of the cascade of molecular events that have occurred. Typically, in unraveling interesting biological phenomena, inferring changes in expression is a critical, yet initial step of the discovery pipeline and the results often feed into downstream interpretive analyses.

Several reports have recently provided snapshots of the performance of gene-level differential expression analysis methods, using both synthetic and experimental data [1, 2, 3, 4, 5]. As an example of the latter, Rapaport et al. [3] used large-scale experimental data from the SEQC (Sequencing Quality Control) consortium, consisting of replicates of well-known cell lines, to evaluate the performance of differential gene expression methods and the effects of modifying sequencing depths and the number of biological replicates. As an extension to this, the SEQC Consortium and Association of Biomolecular Resource Facilities recently finished comprehensive comparison studies with respect to accuracy, sensitivity and reproducibility of *gene-level* measurements across sites, platforms and algorithms [6, 7].

We focus our attention here on a related, important gene expression detection problem: *differential transcript* (or *isoform*) *usage,* or DTU. Previous studies have shown that most multi-exon human genes are affected by alternative splicing and thus can express a variety of different transcripts from the same genomic sequence [8, 9]. Differences in the relative expression of these isoforms between tissues and species are naturally occurring between cell types and allow cells to adapt to the environment, but aberrations from this “normal” splicing pattern can have detrimental consequences for the organism [10]. It is important to distinguish DTU from gene-level differential expression and from transcript-level differential expression (DTE). In particular, DTU considers changes in the *proportions* of the isoforms of a gene that are expressed as opposed to changes of the individual transcript levels. As shown in Figure 1, DTU implies DTE but not necessarily the reverse. Although the main transcriptional units of interest are the transcripts, it has been difficult to obtain accurate and precise transcript-level expression estimates due to the extensive overlap between different transcripts. This has prompted researchers to develop alternative ways of representing and analyzing the observed data. One such approach, which has been used as a surrogate for DTU, is differential exon usage (DEU), where data are represented on the level of disjoint “counting bins”. These bins are transcript “building blocks” similar to exons, and each transcript consists of a combination of counting bins. However, since the counting bins are constructed to be disjoint, bin expression quantification is more straightforward than quantification of the expression of overlapping exons. Preferential inclusion or exclusion of given counting bins points to changes in the expression level of one or more associated transcript(s).

**Figure 1:**
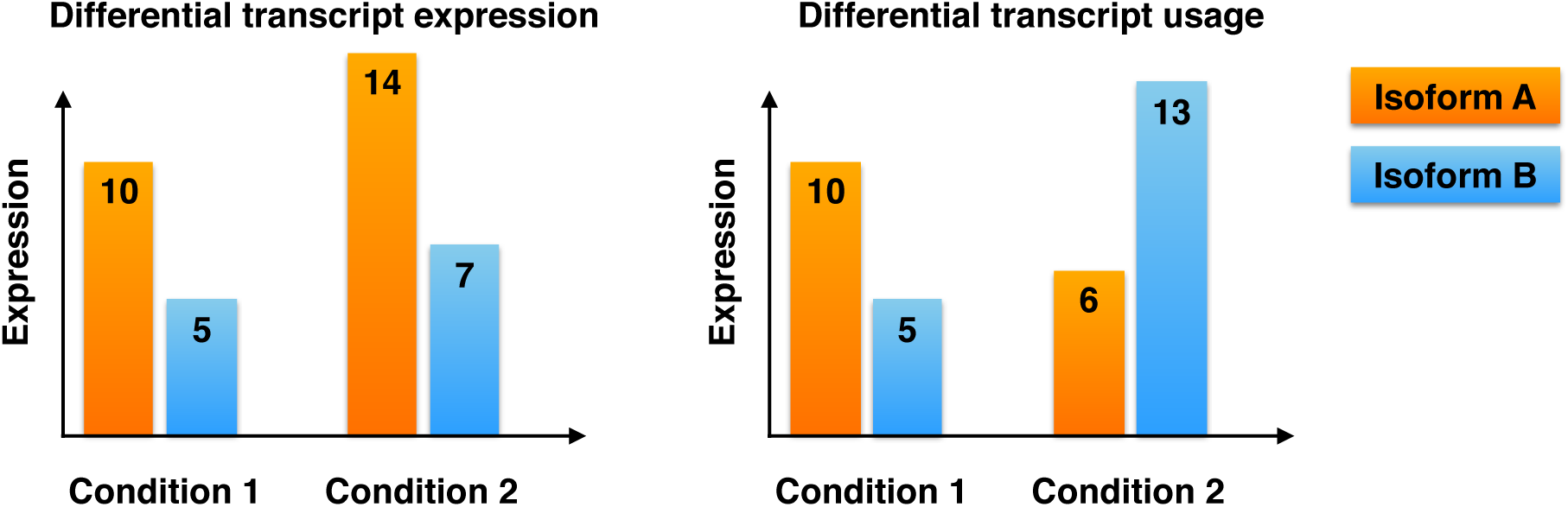
Schematic illustration of differential transcript expression (DTE) and differential transcript usage (DTU) between conditions 1 and 2, for a gene with two isoforms. DTE implies that we can observe expression changes for at least one transcript between condition 1 and condition 2. However, the expression *proportion* of each transcript (as a percentage of the total expression of all transcripts of the same gene) does not necessarily change between conditions, and thus DTE does not necessarily imply DTU. In DTU, on the other hand, the relative expression of the isoforms of a gene changes between the conditions, whereas the total expression of the gene may or may not remain constant. Since at least one isoform must change expression in DTU, it also implies DTE. The numbers indicated on the transcripts in the figure represent expression levels (in arbitrary units) for easier interpretation. Note that in practice, many genes will have more than two isoforms, but the principal difference between DTE and DTU carries over.

A well-placed recent review lists close to 100 computational methods that have been developed in the space of RNA-seq data and splicing analyses [11] and a few comparisons of DTU detection methods have been conducted. For example, a recent software review gives an overview of the features and interfaces available from recent tools to detect DTU, but no critical assessment of empirical performance is given [12]. An early simulation study across a handful of popular mapping-DTU pipelines highlighted that: i) the presence of DTU can increase false positive rates for standard gene-level differential expression analyses, which is perhaps not surprising; ii) in some cases, DTU simply cannot be detected with current methods [13]. In that study, receiver operating characteristic (ROC) curves were presented, but this gives little sense of the FDR control during typical usage. More recently, Liu et. al [14] conducted a wide-ranging comparison of eight DTU detection methods using both simulated and experimental data. They compared performance across different alternative splicing ratios, read depths, dispersion patterns, types of splice changes and sample sizes and explored the influence of annotation, highlighting that no single method dominates in performance and that they often give conflicting results. In the current study, we augment earlier studies in several important ways: we explore a spectrum of counting approaches, highlight striking differences in false discovery rate

(FDR) control between simple and complex organisms, improve existing workflows by applying carefully designed filtering criteria and explore the effect of incomplete annotation catalogs. The simulated data are available from ArrayExpress with accession number E-MTAB-3766.

### Methods for differential transcript usage

Broadly speaking, there are three major classes of methods designed to detect DTU. First, the *assembly-based* (or “isoform deconvolution”) methods (e.g., the cufflinks/cuffdiff pipeline [15, 16, 17]), which reconstruct and quantify the expression of a set of transcripts that best explain the observed reads. The cuffdiff test for differential transcript usage within a gene is based on the Jensen-Shannon divergence, measuring the similarity between two probability distributions. The second class of methods focuses on specific types of alternative splicing (e.g., retained introns or alternative exons) and identifies the number of observed reads that unambiguously support the presence or absence of each splicing event (e.g., rMATS [18]). Comparing these read counts gives an estimate of the “percent spliced in”, or *psi* value, which can then be compared between conditions for each event.

The third type of DTU detection methods do not directly quantify the transcript expression, but rather use differential exon usage as a surrogate to infer DTU. The genome is divided into (typically disjoint) *counting bins* and the number of observed reads overlapping each bin is counted. To infer differential exon (bin) usage between conditions, these methods often make use of (general or generalized) linear models containing an interaction term between the bin identifier and the condition of interest to search for non-proportionality of the bin counts within a gene between the conditions. Arguably the most widely used differential exon usage detection method is implemented in the DEXSeq R package [19] but alternatives, such as the diffSplice function from the limma R package, are available [20]. Since DEXSeq infers differential exon usage, it is left to the user to interpret which transcripts are differentially used, given the evidence for a particular exon bin-condition interaction. However, already knowing which exons are affected can lead to biologically meaningful interpretation of the functional impact of their differential usage (e.g., [19]).

### Bin read counting

The read counting function distributed with the DEXSeq package splits the coding parts of the genes into non-overlapping exon bins and counts the number of reads overlapping each of these bins. Reads that overlap multiple bins are assigned to all of them, which increases the correlation between the bin counts and implies that the sum of the bin counts can significantly exceed the number of sequencing reads in the experiment. With default settings, genes that overlap each other are aggregated into a composite gene complex containing all exon bins of the original genes. The identifier of this complex is obtained by combining the identifiers of the aggregated genes. In practice, this could lead to difficulties in result interpretation, and the differential splicing detection could potentially be affected by overall differential gene expression of a subset of the genes involved in a complex. In this study, we examine whether using alternative definitions of the counting bins, while keeping the interaction inference engine fixed, could provide benefits to the overall performance of DEXSeq. In total, we compare nine variants of counting bins. In this section, we give a brief description of the used methods and define the name that will be used to identify each method in the evaluations (given in parentheses). More details regarding the application of these methods can be found in Additional file 1, as well as in the project GitHub repository (https://github.com/markrobinsonuzh/diff_splice_paper). For more details regarding their theoretical underpinnings and implementation, we refer to the original publications.

In addition to the default DEXSeq exon bin counting *(DEXSeq-default*) (DEXSeq R package v1.14.0) [19], we explore the effect of circumventing the merging of overlapping genes in two ways (see Supplementary Figures 3 and 4 for illustrations). First, we change the arguments to the DEXSeq annotation preparation function to exclude overlapping exon parts on the same strand rather than aggregate genes (*DEXSeq-noaggreg*). This eliminates the composite feature identifiers, but may leave out important genomic regions that can not be unambiguously assigned to one gene. This is particularly problematic if the annotation catalog is “overpopulated” with irrelevant transcripts. Second, we use functions from Bioconductor [21] to “manually” split the genes into disjoint counting bins, excluding overlapping parts even if they are on different strands and count the reads using the `featureCounts` function from the Rsubread R package (v1.18.0) *(featureCounts-flat*) [22, 23], allowing reads to be assigned to multiple features. We also explore the effect of completely circumventing the division of exons into disjoint bins and assign the reads to the original exons, again allowing a read to be assigned to multiple exons (*featureCounts-exon*). The SplicingGraphs R package (v1.8.1) provides a counting method where splicing graphs are constructed from the provided gene models and the reads are assigned to the edges of the graph (representing exons or introns) (*SplicingGraph*). Since the reads spanning exon junctions are expected to be the most informative for identifying DTU, we also explicitly evaluate the use of the junction counts generated by the TopHat aligner (v2.0.14), ignoring all reads that fall completely within exons (*TopHat-junctions*). Using MISO (v0.5.3) [24], we define the counting bins as combinations of isoforms and count the number of reads that align in positions compatible with each given combination (*MISO*). Yet another way of defining counting bins is provided by an intermediate representation from the *casper* R package (v2.2.0) [25], which defines the bins as the exon paths traced out by the two reads in a pair (*casper*). Finally, we define the counting bins as the isoforms themselves and use kallisto (v0.42.1) [26] to estimate the number of reads assigned to each isoform (*kallisto*).

Each of the counting methods generates a count matrix, where the rows correspond to counting bins and the columns to samples. These count matrices are used as the input to DEXSeq in order to search for bins showing evidence of differential usage between conditions. Since each approach defines bins in a different way, and not all bins are possible to unambiguously associate with a given isoform, we evaluate them in terms of their ability to detect differential isoform usage at the gene-level.

Conceptually, each of the different counting bin definitions has advantages and disadvantages when used in conjunction with an inference engine such as DEXSeq, and ultimately the optimal choice likely depends on aspects such as the level at which interpretation can most easily be made, and the confidence in the annotation catalog. DEXSeq allows the user to determine which bin(s) are preferentially included or excluded in a given condition. In some situations, where the differential regulation of specific isoforms are expected to be involved in the phenotype determination, it can be advantageous to define the bins as the isoforms themselves. In other situations, however, the key phenotypic determinant may be the inclusion of a specific exon containing important genetic material, and the precise isoform contributing this exon is less important. In such cases, (sub)exon-specific bins may be preferable. In addition, exon-level bins allows the detection of events (such as exon skipping) that are not annotated as part of an existing isoform, and isoform-level bins will not allow detection of splicing aberrations in genes with a single isoform. On the other hand, exon-level bins could lead to spurious findings in terms of significantly differential bins that could not be the result of a true change in isoform usage.

Also other aspects, such as the degree of “multi-counting”, could lead to performance differences between the bin definition approaches. Typically, transcript abundance estimation methods, such as kallisto, attempt to distribute reads overlapping multiple transcripts in order to optimize a likelihood function. In contrast, the DEXSeq counting script assigns reads overlapping multiple bins to each of them, potentially increasing the correlation between bin counts within a gene. This multi-counting increases the count for the individual bins, particularly in situations where the bins are much shorter than the reads, and thus potentially leads to higher statistical power. On the opposite side of the spectrum, methods that consider only junction-spanning reads (such as *TopHat-junctions*) excludes a potentially large fraction of the reads and can thus be expected to lose power, especially when the exons are relatively long compared to the reads.

For methods based on the original (non-disjoint) exons (such as *featureCounts-exon*), we expect a lower power to detect switches between isoforms where the critical region (the genomic region that is unique to one or the other isoform) is small. The reason for this is that the reads from the critical region will contribute relatively little to the counts of the exons (bins). Thus, even dramatic relative changes in this small contribution may pass unnoticed, whereas it would be apparent if the critical region formed a bin in itself (such as, for example, with *DEXSeq-default, DEXSeq-noaggreg* and *featureCounts-flat*).

## Results

We evaluated the counting methods outlined in the previous section (using DEXSeq as the inference engine) as well as cuffdiff (v2.2.1) and rMATS (v3.0.9) using simulated data based on fruit fly and human as described in the Methods section. The characteristics of the two organisms are outlined in Additional file 1:Supplementary Figure 1 and Supplementary Table 1. For each organism, we simulated reads for 3 replicates in each of two conditions. DTU was introduced for 1,000 genes by reversing the relative abundances of the two most abundant isoforms in one of the conditions, while keeping the total number of transcripts generated from the gene constant (that is, no gene-level differential expression). This generated truly differential genes with a range of “effect sizes” (the difference in relative abundance between the two differentially used isoforms). The simulation framework is outlined in Additional file 1:Supplementary Figure 2. In Additional file 1:Supplementary Figures 7-8, we also provide a comparison of the overall count patterns for the different methods.

**Figure 2:**
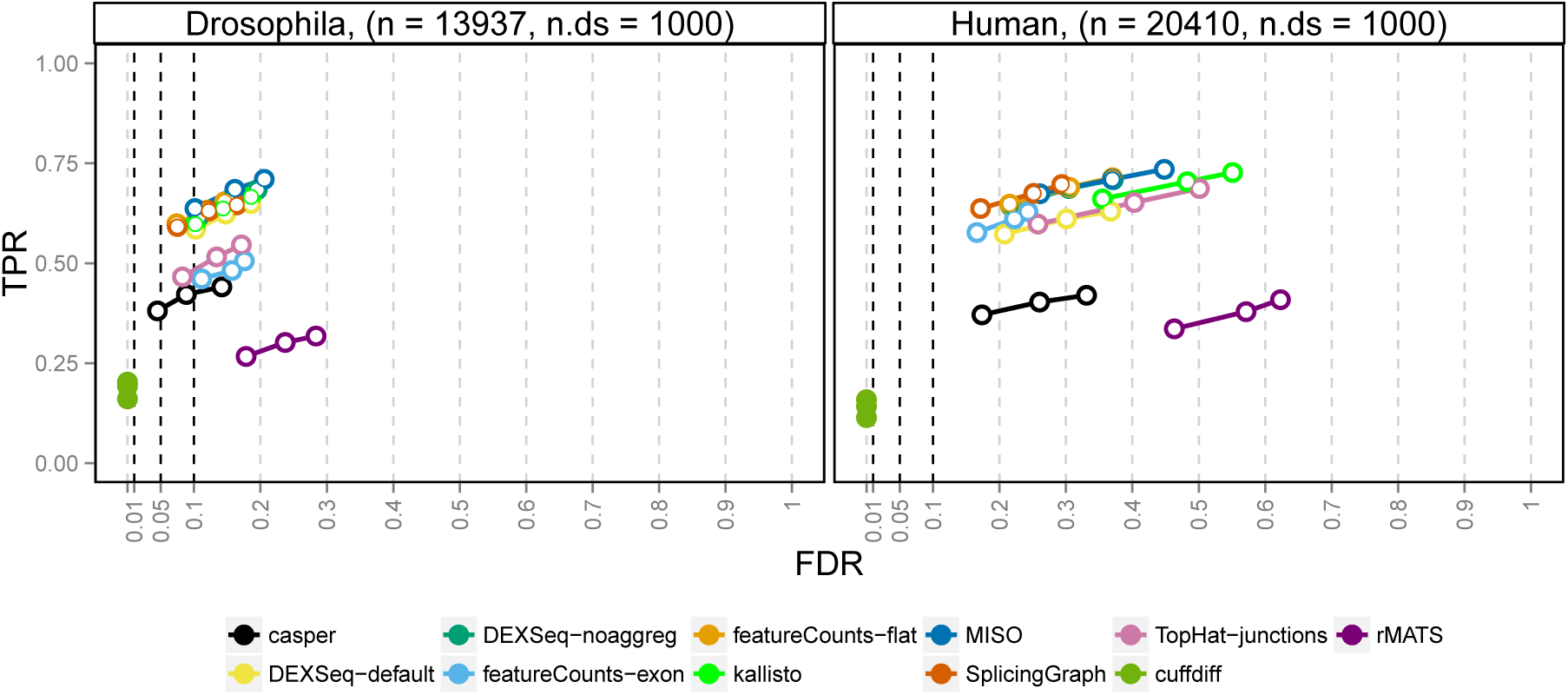
Overall performance of the evaluated methods. The three circles for each method indicate the observed false discovery rate (FDR) and true positive rate (TPR) when the gene-wise q-values are thresholded at three commonly used thresholds: 0.01, 0.05 and 0.1. Ideally each circle should fall to the left of the corresponding vertical line, since this would indicate that the FDR is controlled at the imposed level. A circle is filled if the FDR is controlled and open otherwise. The total number of genes (n) as well as the number affected by differential isoform usage (n.ds) are given in the panel headers. Overall, only *cuffdiff* manages to control the FDR, but at the cost of a reduction in power (TPR). The FDR control is worse in the human simulation than in the fruit fly simulation, potentially due to the larger number of isoforms in the human transcriptome.

**Table 1:**
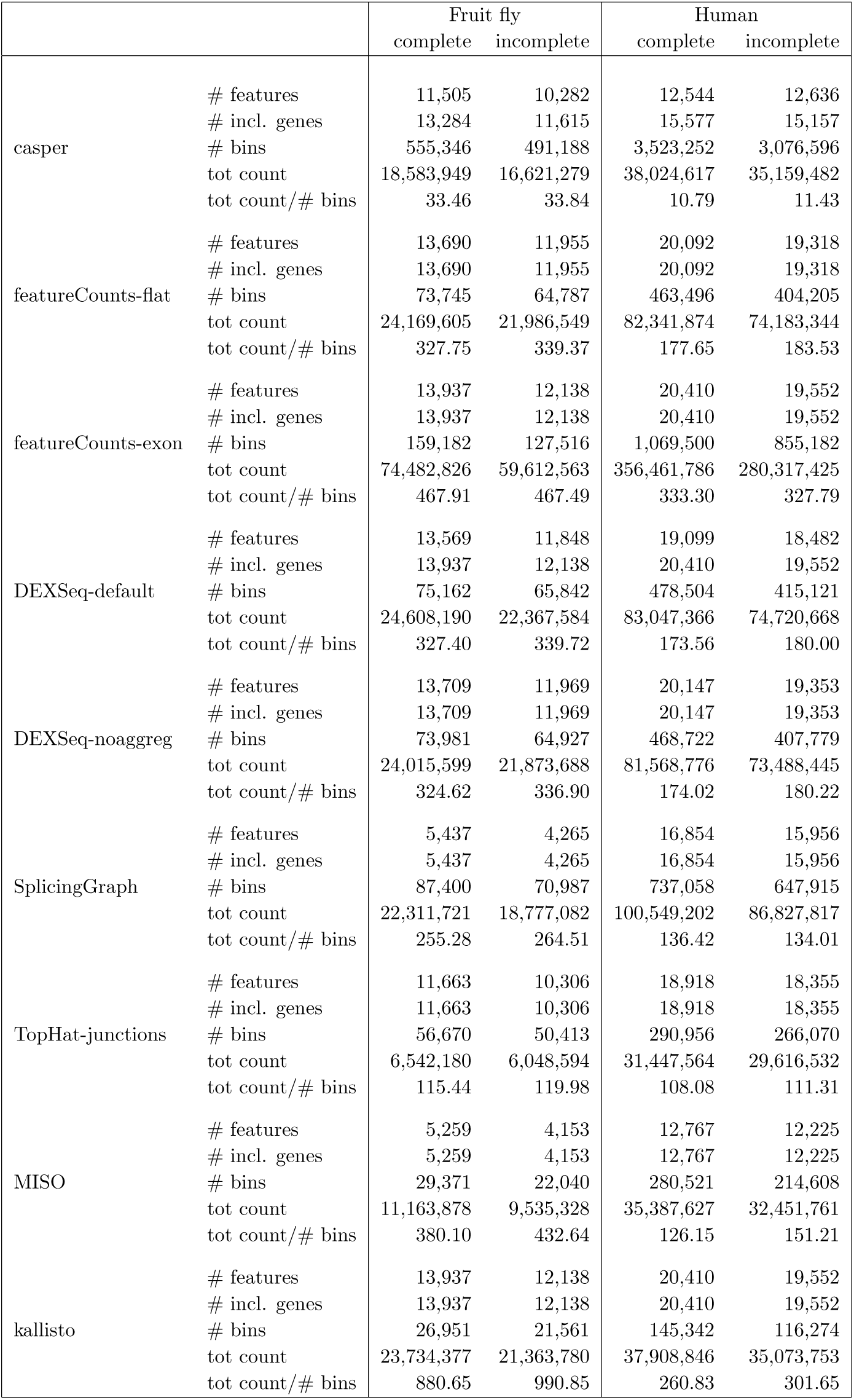
The number of features (genes + gene complexes), the number of genes covered by any of the features (# incl. genes), the number of counting bins and the sum of all bin counts for various counting methods and for the complete and reduced annotation catalog. Randomly excluding 20% of the transcripts from the annotation catalog generally led to a slight increase in the average bin count, with the most pronounced effects seen for the methods defining the counting bins as (combinations of) transcripts (*kallisto* and *MISO*). *MISO* and *SplicingGraph* consider only genes with at least two isoforms, which explains the lower number of features for these methods.

|  |  | Fruit fly |  | Human |  |
| --- | --- | --- | --- | --- | --- |
|  |  | complete | incomplete | complete | incomplete |
| casper | # features | 11,505 | 10,282 | 12,544 | 12,636 |
|  | # incl. genes | 13,284 | 11,615 | 15,577 | 15,157 |
|  | # bins | 555,346 | 491,188 | 3,523,252 | 3,076,596 |
|  | tot count | 18,583,949 | 16,621,279 | 38,024,617 | 35,159,482 |
|  | tot count/# bins | 33.46 | 33.84 | 10.79 | 11.43 |
| featureCounts-flat | # features | 13,690 | 11,955 | 20,092 | 19,318 |
|  | # incl. genes | 13,690 | 11,955 | 20,092 | 19,318 |
|  | # bins | 73,745 | 64,787 | 463,496 | 404,205 |
|  | tot count | 24,169,605 | 21,986,549 | 82,341,874 | 74,183,344 |
|  | tot count/# bins | 327.75 | 339.37 | 177.65 | 183.53 |
| featureCounts-exon | # features | 13,937 | 12,138 | 20,410 | 19,552 |
|  | # incl. genes | 13,937 | 12,138 | 20,410 | 19,552 |
|  | # bins | 159,182 | 127,516 | 1,069,500 | 855,182 |
|  | tot count | 74,482,826 | 59,612,563 | 356,461,786 | 280,317,425 |
|  | tot count/# bins | 467.91 | 467.49 | 333.30 | 327.79 |
| DEXSeq-default | # features | 13,569 | 11,848 | 19,099 | 18,482 |
|  | # incl. genes | 13,937 | 12,138 | 20,410 | 19,552 |
|  | # bins | 75,162 | 65,842 | 478,504 | 415,121 |
|  | tot count | 24,608,190 | 22,367,584 | 83,047,366 | 74,720,668 |
|  | tot count/# bins | 327.40 | 339.72 | 173.56 | 180.00 |
| DEXSeq-noaggreg | # features | 13,709 | 11,969 | 20,147 | 19,353 |
|  | # incl. genes | 13,709 | 11,969 | 20,147 | 19,353 |
|  | # bins | 73,981 | 64,927 | 468,722 | 407,779 |
|  | tot count | 24,015,599 | 21,873,688 | 81,568,776 | 73,488,445 |
|  | tot count/# bins | 324.62 | 336.90 | 174.02 | 180.22 |
| SplicingGraph | # features | 5,437 | 4,265 | 16,854 | 15,956 |
|  | # incl. genes | 5,437 | 4,265 | 16,854 | 15,956 |
|  | # bins | 87,400 | 70,987 | 737,058 | 647,915 |
|  | tot count | 22,311,721 | 18,777,082 | 100,549,202 | 86,827,817 |
|  | tot count/# bins | 255.28 | 264.51 | 136.42 | 134.01 |
| TopHat-junctions | # features | 11,663 | 10,306 | 18,918 | 18,355 |
|  | # incl. genes | 11,663 | 10,306 | 18,918 | 18,355 |
|  | # bins | 56,670 | 50,413 | 290,956 | 266,070 |
|  | tot count | 6,542,180 | 6,048,594 | 31,447,564 | 29,616,532 |
|  | tot count/# bins | 115.44 | 119.98 | 108.08 | 111.31 |
| MISO | # features | 5,259 | 4,153 | 12,767 | 12,225 |
|  | # incl. genes | 5,259 | 4,153 | 12,767 | 12,225 |
|  | # bins | 29,371 | 22,040 | 280,521 | 214,608 |
|  | tot count | 11,163,878 | 9,535,328 | 35,387,627 | 32,451,761 |
|  | tot count/# bins | 380.10 | 432.64 | 126.15 | 151.21 |
| kallisto | # features | 13,937 | 12,138 | 20,410 | 19,552 |
|  | # incl. genes | 13,937 | 12,138 | 20,410 | 19,552 |
|  | # bins | 26,951 | 21,561 | 145,342 | 116,274 |
|  | tot count | 23,734,377 | 21,363,780 | 37,908,846 | 35,073,753 |
|  | tot count/# bins | 880.65 | 990.85 | 260.83 | 301.65 |

The performance evaluation is based on gene-level q-values (adjusted p-values) calculated as described in the Methods section. After thresholding the gene-wise q-values at three common values (0.01, 0.05 and 0.1), we evaluated the true positive rate (TPR, the fraction of genes with true differential isoform usage that show q-values below the threshold) and the observed false discovery rate (FDR, the fraction of all genes with q-value below the threshold where there is no true differential isoform usage). Any gene identifiers that were not present in the list of simulated genes were excluded from the evaluation, since these genes could not be classified as either true positive, true negative, false positive or false negative. This affected the performance estimates for *casper* and *DEXSeq-default* counts, as well as *cuffdiff*, since these methods sometimes form gene complexes, with identifiers that are combinations of the identifiers of the merged genes. The number of features (genes + complexes) and the number of complexes for these methods are given in Additional file 1:Supplementary Table 1.

### Overall performance and effect of transcriptome complexity

Overall, the observed TPRs were similar between the two studied organisms, while the achieved FDRs were higher in the human simulation (Figure 2). We hypothesized that the difference in performance between the fruit fly and human simulation was associated with the higher level of complexity of the human transcriptome, with many more isoforms per gene. Indeed, stratifying the results by the number of isoforms for each gene showed a clear association between the number of isoforms and the observed FDR in both organisms (Figure 3). On a global scale, only *cuffdiff* managed to satisfactorily control the FDR in any of the organisms, but at the price of much lower power than most other methods. The conservative nature of *cuffdiff* has been previously described in the context of differential transcript expression [27]. One major reason for the conservativeness in the current study is that only genes where the coding output differs between the differentially used isoforms can be detected. Applying *cuffdiff* with an artificial annotation file where CDS and protein ID entries were replaced by exon and transcript IDs, respectively, considerably improved the power of *cuffdiff* (Additional file 1:Supplementary Figures 18-19). Another contributing reason to the low power could be that the sampling scheme used by *cuffdiff* to evaluate significance currently does not yield nominal p-values below 5 · 10^−5^.

**Figure 3:**
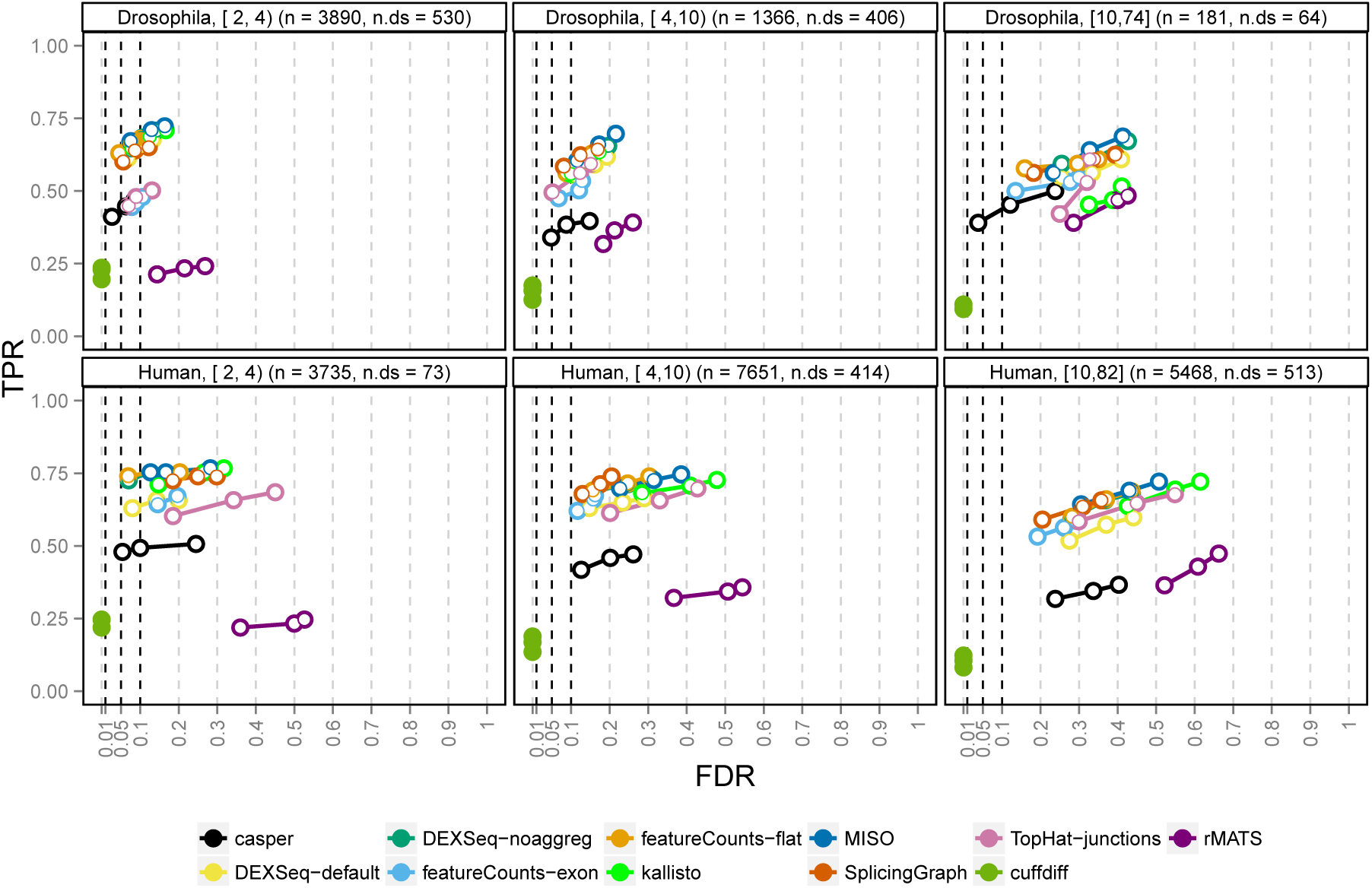
Performance stratified by number of isoforms per gene. The ability to control the false discovery rate (FDR) at an imposed level depends on the number of isoforms of the genes (indicated in the panel headers as e.g. [2, 4)). The FDR control for genes with many isoforms is worse than that for genes with few isoforms. The total number of genes (n) as well as the number of genes affected by DTU (n.ds) in each category are indicated in the panel headers.

The second lowest power was obtained by *rMATS*, mostly since it is only able to detect simple splicing events (for these, however, the power was comparable to the other methods, see Additional file 1:Supplementary Figure 12). Among the counting methods used with DEXSeq, the exon path counts from *casper* showed the lowest power for both organisms. This could be attributed to a combination of the relatively low read count for the individual bins and the merging of features into complexes, which could not be directly matched with the list of truly differential or non-differential genes. The same merging effect was seen for *cuffdiff* and for *DEXSeq-default* (which therefore showed a lower TPR than *DEXSeq-noaggreg*), albeit not as strong since *DEXSeq* and *cuffdiff* only merge genes if they are on the same strand, while *casper* merges also overlapping genes on different strands. The merging effect was particularly pronounced in the human data, where a larger fraction of the genes are overlapping to some extent and thereby were aggregated into complexes (Additional file 1:Supplementary Table 1). An alternative could be to evaluate the methods only based on the genes for which a true status indicator as well as a q-value were available (that is, to subset the truth and result tables to only these genes before the performance evaluation). This approach is explored in Additional file 1 (Supplementary Figure 17). As expected, it improved the power of *casper* and *DEXSeq-default* substantially.

In the fruit fly simulation, the count matrices generated by *featureCounts-exon* and *TopHat-junctions* performed relatively poorly, while their results in the human simulation were more on par with the other methods. The performance difference between the organisms may be associated with the relatively low junction counts in fruit fly and with the observed inability of *featureCounts-exon* to detect switches between genetically similar isoforms, where the critical regions (the regions that are unique to one of the isoforms) are very small (see Additional file 1:Supplementary Figure 11 for an illustration). In both organisms, the *SplicingGraph, DEXSeq-noaggreg* and *featureCounts-flat* counts performed similarly. This was not surprising since the counting bins were similar between these methods (exons and introns). The transcript-level counts obtained by *kallisto* also appeared to give good results when combined with the DEXSeq inference engine, especially for relatively simple genomes such as the fruit fly. Similarly, *MISO* counts provided good results in the fruit fly simulation, but showed a relatively high FDR for the human data. In addition to its low power due to the inability to detect complex splicing events, *rMATS* showed a higher FDR than the other methods. A closer examination of the false positive events revealed that these are enriched with rare events (inclusion levels close to 0) and relatively depleted of events with inclusion levels in the middle range between 0 and 1 (see Additional file 1:Supplementary Figure 20).

### Performance stratified by gene characteristics

To better understand the results shown in Figure 2, we evaluated the performance within given gene strata, defined by various annotations. In Figure 4, we show the effect of stratifying by the difference between the relative abundances of the two most highly abundant isoforms (which are also the ones that are differentially used for the genes affected by DTU). Additional file 1 (Supplementary Figures 9 and 11-12) contains similar figures stratified by other gene annotations, such as overall gene expression level, compositional similarity between the differentially used isoforms and the type of alternative splicing event. Differential isoform usage was introduced by reversing the two most abundant isoforms for the 1,000 genes selected and as expected, the larger this difference, the easier we could detect alternative usage (Figure 4). The performance of all methods, for both organisms, increased dramatically between the group of genes where the relative abundance difference was below 33% and the group with relative abundance differences between 33% and 67%. For the genes with one highly dominant transcript (relative abundance difference above 67%), although almost all truly differential genes were found, the FDR control was very poor, especially in the human simulation.

**Figure 4:**
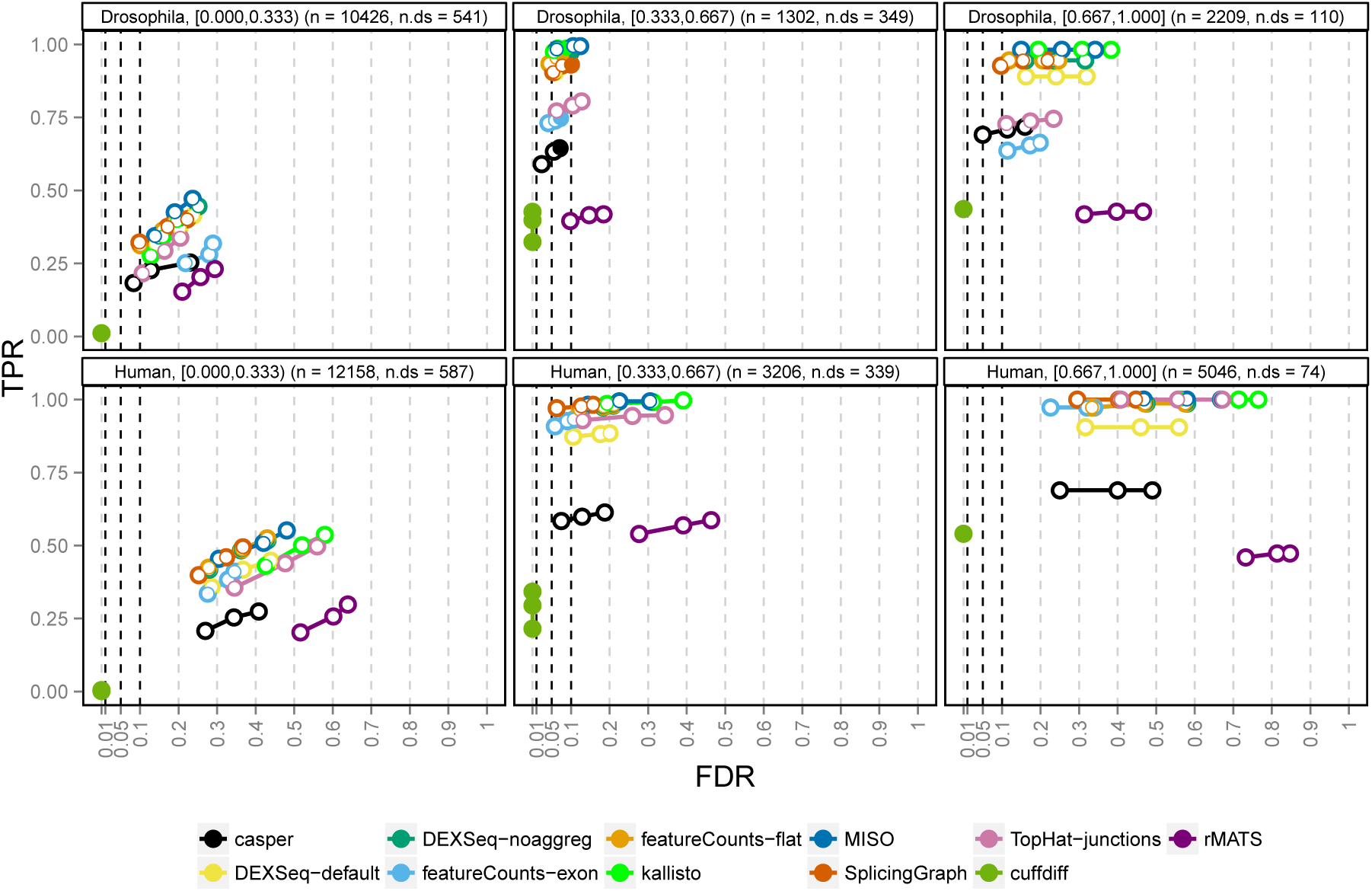
Performance stratified by isoform dominance. The degree of isoform dominance (the difference in relative abundance between the two most highly expressed isoforms of a gene, indicated in the panel headers as e.g. [0.000,0.333)) is the main determinant of the methods’ ability to detect differential isoform usage in our simulations (recall that the differential isoform usage was introduced by switching the abundances of the two most dominant transcripts of 1,000 selected genes). DTU for genes where the two differentially used isoforms are expressed at almost the same level (left panels) is more difficult to detect than genes with one dominant isoform (right panels). On the other hand, the FDR control in the latter category is very poor for most methods. The total number of genes (n) as well as the number of genes affected by DTU (n.ds) in each category are indicated in the panel headers.

### Effect of annotation catalog incompleteness on counting methods

All methods evaluated in this study rely on known annotations. While both organisms chosen here have well-characterized transcriptomes, this is far from true for many non-model species. Here, we studied the effect of an incomplete annotation catalog on the power of detecting differential isoform usage, by excluding 20% of the known transcripts from the reference gtf file (randomly chosen, but proportionally split between differentially used and non-differentially used). We then re-ran the analysis of the simulated data, starting from the TopHat read alignment step. We restricted this analysis to the counting methods combined with DEXSeq, since they provided the best performance above.

Excluding isoforms led to a reduction in the number of counting bins for all methods (Table 1) and for most of the methods a slight increase in the average read count per bin. The methods assigning reads to (combinations of) transcripts (*MISO* and *kallisto*) showed the largest relative increase in the average read count per bin. The incomplete annotation catalog had a much greater impact on the detection power of the various counting methods in the fruit fly simulation than in the human simulation (Figure 5). A likely explanation for this effect is the low number of isoforms per gene for the fruit fly, implying that there will be many genes with only one (or even zero) isoform remaining in the annotation catalog after the exclusion. Indeed, stratifying the results by the number of excluded differentially used isoforms and the total number of retained isoforms showed that the number of remaining isoforms was the strongest determinant of the performance (Additional file 1:Supplementary Figures 13-16). Many of the counting methods were able to detect the differential isoform usage even after excluding both the differentially used isoforms from the annotation, as long as the total number of remaining isoforms was large enough. The methods using the transcript structure to define the counting bins (*kallisto, MISO* and *SplicingGraph*) were most strongly negatively affected when only one isoform was retained. This can be attributed to the default exclusion of genes with a single isoform by *SplicingGraph* and *MISO,* and that *kallisto* only generates a single counting bin for these genes. Conversely, *featureCounts-exon* and *TopHat-junctions* were least affected.

**Figure 5:**
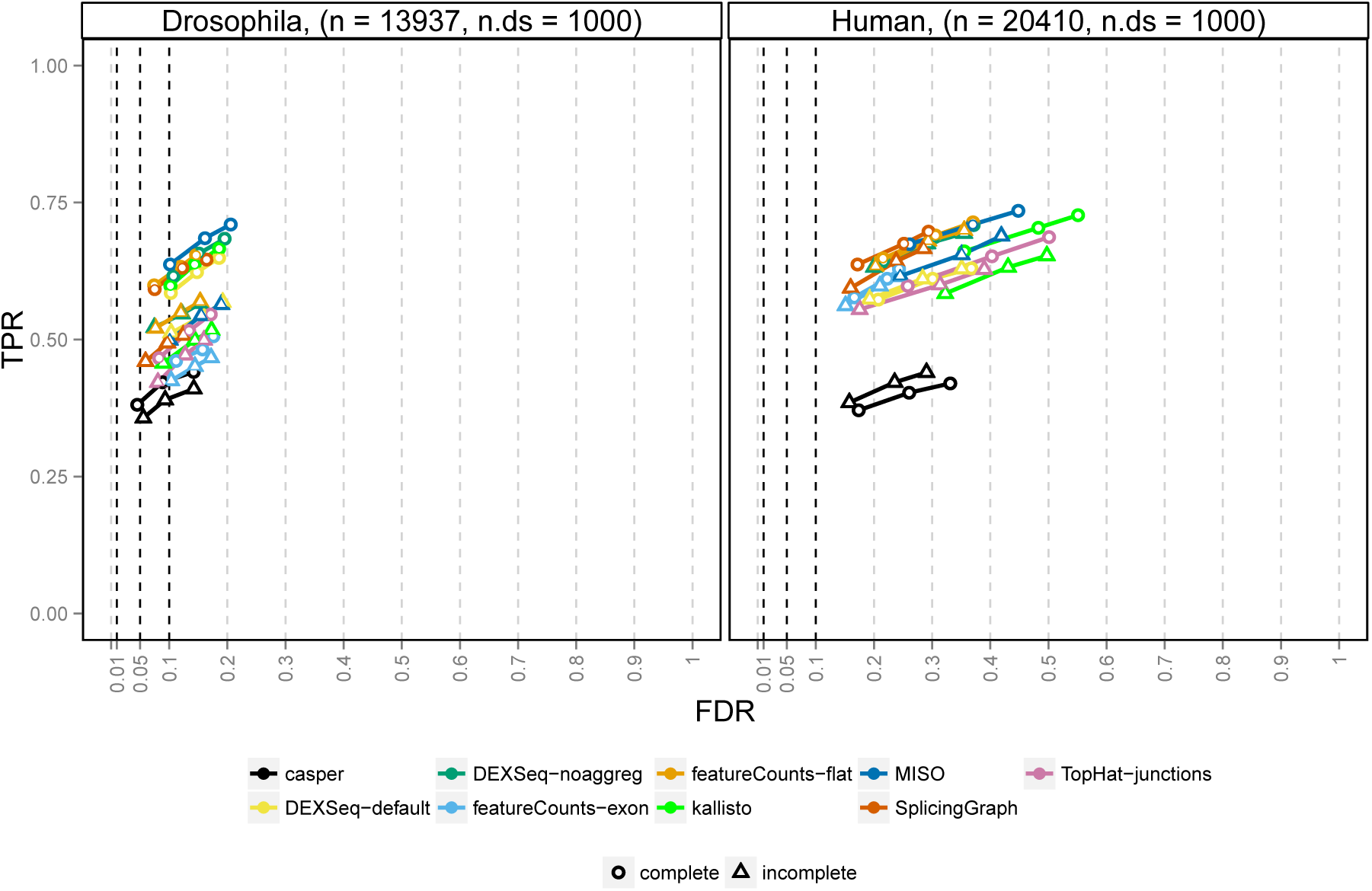
TPR and FDR stratified by incompleteness of annotation. The circles indicate the performance of the different methods using the complete annotation catalog. The triangles indicate the performance achieved when the analysis was performed using an incomplete annotation catalog, where 20% of the transcripts were excluded. The overall impact of the incompleteness of the annotation catalog is larger for the fruit fly simulation (left panel) than for the human simulation (right panel). This appears to be mainly due to the larger number of isoforms in the human transcriptome and the associated larger redundancy. Additional file 1:Supplementary Figures 13-16 stratify the results further and provide additional insight.

### Isoform pre-filtering improves the observed FDR

We noted above that the empirical FDR was often much higher than the imposed FDR threshold, especially for genes with one largely dominating isoform and that this effect was consistent across most counting methods. In an attempt to better understand the reasons behind this high FDR, we compared the characteristics of the genes falsely called differentially spliced (the false positives, FP) to those of the true positives (TP, correctly called genes with true DTU), the false negatives (FN, non-significant genes with true underlying DTU) and true negatives (TN, genes correctly called non-differential). We focused on the counts obtained by *DEXSeq-noaggreg*, due to their intrinsic link to the DEXSeq framework and their overall good performance in the previous evaluations. The most notable observation was that the FP genes showed a larger variance among the dispersion estimates for their respective exon bins than the other gene categories (Additional file 1:Supplementary Figures 21-22). The effect was more pronounced for genes with many counting bins and genes with one highly dominant isoform. We hypothesized that reducing the number of bins by an initial filtering step, eliminating the non-expressed isoforms from the annotation catalog used to generate the counting bins, could improve the FDR control. Based on the RSEM-estimated isoform percentages from which the individual sample isoform percentages were derived, we thus generated four new annotation files for the DEXSeq counting by excluding all isoforms with relative abundance below (in turn) 5%, 10%, 15% and 25% in both conditions. Then, we performed the counting and differential exon usage testing as before. Other filtering paradigms could be imagined, such as successively excluding the lowest expressed isoforms as long as their total relative abundance does not exceed a fixed threshold. For all the thresholds we evaluated, the reduction in the number of bins was substantially larger than the relative reduction in total bin counts (Additional file 1:Supplementary Tables 2-3), which suggests a high degree of redundancy and/or that few reads were assigned to the excluded bins. The human simulated data contained a larger fraction of non-expressed isoforms (Additional file 1:Supplementary Table 1) and thus the reduction in the number of retained isoforms was larger than for the fruit fly simulation. It is worth highlighting that isoform-level filtering is not equivalent to filtering bins out of the count table (see Additional file 1 for a comparison to bin filtering approaches).

Excluding lowly abundant isoforms from the catalog before forming the counting bins provided a substantial performance improvement compared to unfiltered data (Figure 6), in particular for the previously problematic genes with one largely dominant transcript. Already excluding isoforms with relative abundance less than 5% provided a tangible improvement in FDR control. Comparing the FPs found with the original annotation file to those found after filtering at the 5% level revealed that in the human simulation, 184 genes were called FP exclusively with the original file, while 30 were called FP exclusively after filtering. Studying these genes in more detail revealed that many of the 30 FPs exclusive to the filtered setting were likely significant mainly due to a less stringent correction for multiple comparisons (fewer counting bins) and showed evidence of differential usage also with the original annotation (Additional file 1:Supplementary Figures 23-25). Conversely, the 184 genes called FP exclusively with the original annotation were correctly classified after the filtering mainly since the bins contributing to the significance of the gene were filtered out (Additional file 1:Supplementary Figures 26-28). Notably, the strategy of excluding lowly expressed transcripts is of general utility, since it also significantly reduces the rate of false discoveries for DEXSeq combined with *kallisto* estimated counts (Additional file 1:Supplementary Figures 34-35). Likewise, setting a higher threshold on the absolute change in percent-spliced-in reduced the false discoveries called by *rMATS* with a modest drop in power (Additional file 1:Supplementary Figures 36).

**Figure 6:**
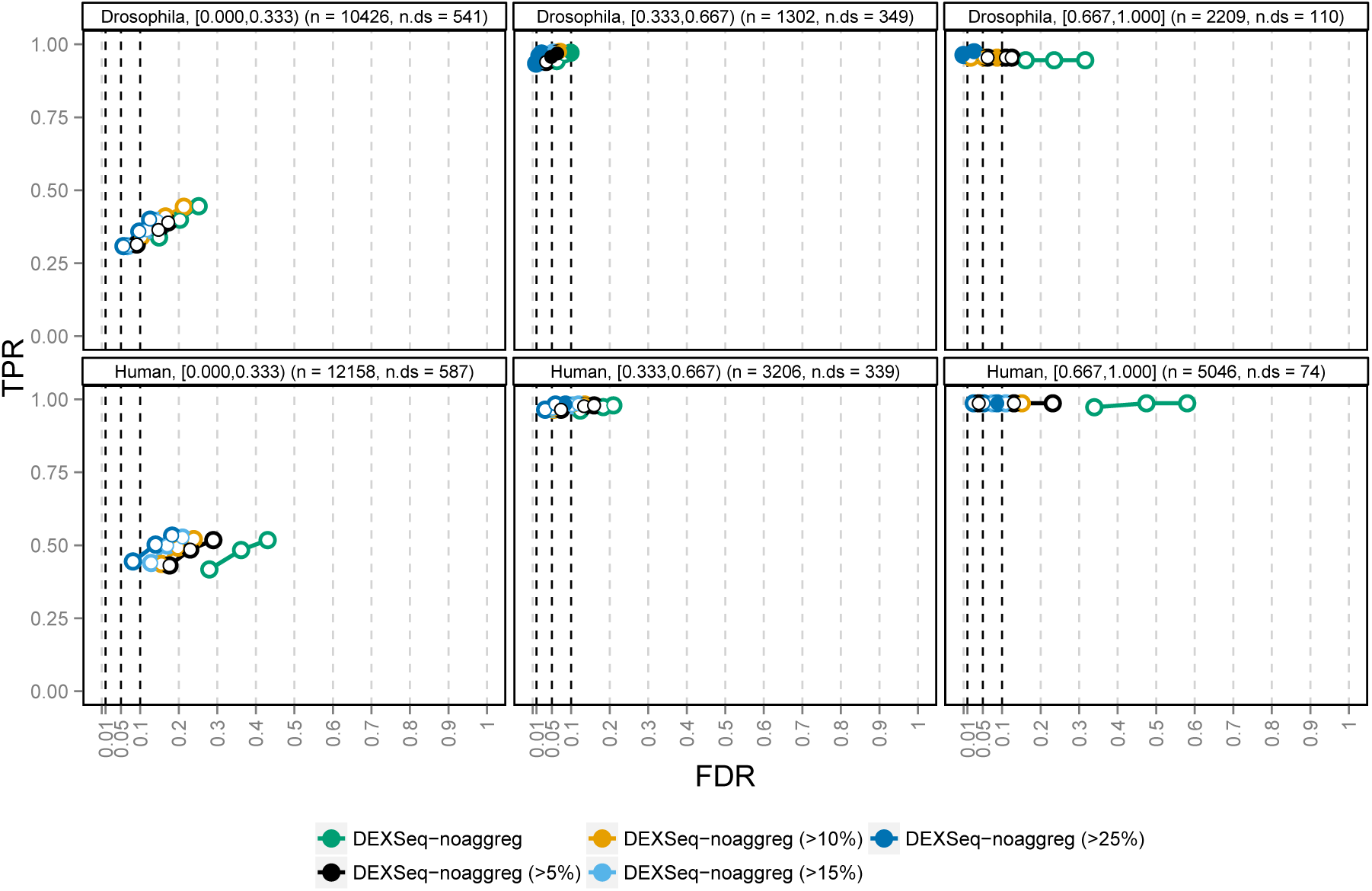
TPR and FDR for *DEXSeq-noaggreg,* with various degrees of isoform-guided annotation filtering. The performance evaluation is stratified according to the degree of isoform dominance (the difference in relative abundance between the two most highly expressed isoforms of a gene, indicated in the panel headers as e.g. [0.000,0.333)). Excluding lowly expressed transcripts from the annotation catalog before defining the *DEXSeq-noaggreg* counting bins improves the control of FDR, with negligible loss (or even a slight gain) of power. The total number of genes (n) as well as the number of genes affected by DTU (n.ds) in each category are indicated in the panel headers.

In our simulation study, we were able to use the underlying isoform abundances to perform the filtering. Of course, in experimental data sets, the true relative isoform abundances are not known and they may vary between samples and conditions. A possible approach in a real-data setting could be to first estimate the relative isoform abundance using, e.g., kallisto [26] or RSEM [28] and exclude isoforms where the relative abundance estimate does not exceed a certain fraction in any of the studied samples. This approach is evaluated in Additional file 1, along with several other filtering paradigms based on the observed exon bin counts (Additional file 1:Supplementary Figures 29-33). There, we show that the isoform-guided filtering, based on *estimated* abundances, performs equally well. Another potential complication in experimental data is that also lowly abundant isoforms can be subject to differential usage and so already a low filtering threshold may exclude differentially used isoforms. However, depending on the experiment, the reduction in FDR may compensate for this loss.

## Discussion and Conclusions

In this study, we have used simulated data to evaluate nine counting methods in terms of their compatibility with the DEXSeq inference engine for detection of differential transcript usage. The main motivation was to evaluate whether a non-standard counting approach could improve the performance of a well-established statistical inference method and to what extent the counting protocol influences the performance. Importantly, we did not attempt to quantify or compare the “correctness” of the counts obtained by the different methods, or their potential compatibility with other inference frameworks. We also compared the counting methods to one assembly-based method *(cuffdiff)* and one event-based method *(rMATS).* Our results show that the inference framework provided by DEXSeq, testing for the differential inclusion of particular counting bins, works well in conjunction with many different bin definitions, from sub-exonic bins quantified using data aligned to the genome (e.g., *DEXSeq-noaggreg*) to full-length transcript bins quantified using alignment-free approaches (e.g., *kallisto). DEXSeq-noaggreg* has the distinct advantage that some types of *unannotated* events (e.g., exon skipping) can still be detected with default settings, whereas *kallisto* has a definite speed advantage. We expect that despite the recent excitement around ultrafast alignment-free quantification methods, many researchers will still elect to place reads in genomic context using alignment-based methods for visualization and interpretation. Ultimately, the choice of bin type should be guided by the level (exon, transcript) at which relevant inference about the underlying biology can be made.

The characteristic that most strongly influenced the power to detect differential isoform usage was, not surprisingly, the magnitude of the change in proportions between the differentially used isoforms. Also the complexity of the gene, that is, the number of isoforms, had a noticeable impact on the specificity and many complex but truly non-differentially spliced genes were falsely considered differential. Taken together, the count-based methods showed strong performance compared to the other methods. The best performance was obtained with the exon bin counts from *DEXSeq-noaggreg* (that, is the standard DEXSeq counting pipeline but without the default aggregation of overlapping genes) and *featureCounts-flat.* These methods provided a solid performance independently of the alternative splicing event type, the degree of similarity between the differentially used isoforms, the overall expression level and the number of isoforms for the genes and the distribution of relative isoform abundances. The default DEXSeq counting, which aggregates overlapping genes into complexes, showed worse performance than counting without the aggregation step. This was mostly due to problems matching the identifiers of the detected differentially spliced genes (complexes) with the list of truly differentially spliced ones, which could also be a problem when interpreting results from an experimental data set.

*SplicingGraph, MISO* and *kallisto,* which explicitly make use of the transcript structure to generate the counting bins, worked as well as the exon bin counts as long as the annotation catalog was complete, but the default exclusion of genes with a single annotated isoform *(SplicingGraph* and *MISO),* or the generation of only a single counting bin for those genes *(kallisto)* was detrimental in the situation with a relatively simple transcriptome (fruit fly) and an incomplete transcript catalog. The exon path counts from *casper* showed low true positive rates, due to a combination of the low counts for each path and an aggregation of identifiers for overlapping genes similar to the one by *DEXSeq-default*. *TopHat-junctions* counts showed poor performance in the fruit fly simulation, especially for genes with few isoforms, which may be due to the relatively long fruit fly exons, implying a low number of reads spanning exon junctions. Using featureCounts on the original exon level (that is, without “flattening” the reference annotation) gave poor results when the counts were analyzed with DEXSeq, especially for the fruit fly simulation. These counts were not able to capture differential isoform usage when the differentially used isoforms were similar and were generally unable to detect most common simple alternative splicing event types.

By comparing the results from the human and fruit fly simulations, we have shown that the performance of the counting methods as well as the inference framework in general is dependent on characteristics and annotation completeness of the underlying transcriptome. The higher complexity of the human transcriptome led to a higher overall false discovery rate than for the fruit fly simulation, especially for genes with one dominant transcript. In the fruit fly genome, with long exons and few transcripts per gene, incomplete annotation had a detrimental effect on the power to detect differentially used isoforms while in the human simulation, where the exons are shorter and the number of transcripts per gene is much larger, the redundancy dampened the effect of missing annotations considerably. The negative impact of missing annotation entries could potentially be remedied by extending the annotation catalog using tools such as cufflinks [29], although it may not always be unambiguously clear if a newly assembled isoform represents a variant of a gene already existing in the annotation (and if so, which one) or if it should be considered separately.

Finally, we showed that the high false discovery rate obtained especially for genes with one largely dominant isoform could be substantially improved by a simple pre-filtering of the isoform catalog before the construction of the counting bins, with negligible loss of power. Excluding lowly abundant isoforms in this way led to a tangible reduction in the number of counting bins, with only a small reduction in the number of reads assigned to the genes. This type of filtering can also be biologically justified, given previous studies that have suggested that “noisy” or erroneous splicing is responsible for the majority of very low-abundance isoforms [30, 31]. Our evaluations suggest that the proposed isoform-guided filtering leads to better results than filtering of counting bins after the counting, which is the current standard procedure.

The analyses in this study are based on simulated data, due to the lack of publicly available, extensively validated experimental data sets for splicing analyses. By using experimental data to estimate the transcript abundances underlying the simulation, we capture aspects such as the number and relative expression of isoforms within each gene, and the correlation structure of expression levels between genes. The choice of RSEM as the simulator of transcript expression was motivated by its speed, ease of use and flexibility, as well as the ability to estimate and incorporate sequencing biases and error structure from experimental data. However, experiments with other simulation engines (not shown) gave very similar results and we do not expect the choice of simulator to have a major impact on the presented results. All comparisons between methods are performed on the gene level, since this is the “least common denominator” of the different bin definition approaches and since, as discussed in the introduction, the choice of bin definition in a real scenario depends largely on the level at which relevant interpretation can be made. It it also worth noting that differential transcript usage (DTU), which is the topic of this paper, could affect also transcripts that are not themselves differentially expressed, since it focuses on the *relative* abundance of a gene’s isoforms.

The simulated data we used is made available in standard formats with accompanying metadata (see ’Availability of supporting data’ below), and thus our performance benchmark can be readily extended as new method innovations are made. The Supplementary website also gives a link to a web application that facilitates the recreation of the performance comparisons shown in our study.

## Methods

### Simulation

This section describes in detail the simulation procedure employed to generate the fastq files that are the basis for the evaluation of the various methods (see Additional file 1:Supplementary Figure 2 for a schematic illustration). We performed one simulation for human and one for fruit fly, following the same principles. For each organism, we simulated data from two conditions, each with three biological replicates. These two organisms were chosen since their transcriptomes show different characteristics, which could potentially have a large impact on the performance of the evaluated methods (Additional file 1:Supplementary Figure 1 and Supplementary Table 1). Most strikingly, fruit fly has considerably longer exons and transcripts than the human, while human genes typically have a much larger number of isoforms per gene. Moreover, they are model organisms for which the transcriptome catalog can be expected to be at least reasonably well characterized and, as such, the results translate to many real studies. However, we emphasize that the results are not expected to be restricted to these organisms, and we make our code available to facilitate extensions to other organisms.

### Reference files

The simulation was performed using RSEM (v1.2.21) [28] which, given TPM (transcripts per million) expression values for each isoform in a given reference annotation catalog, simulates the sequencing process and generates fastq files. In order to get realistic values for the expression levels and relative isoform abundances, as well as for the sequencing parameters, we used RSEM to estimate these values from real RNA-seq data sets (by means of the `rsem-calculate-expression` module). For the human simulation, we used the fastq files from the sample SRR493366 (http://www.ebi.ac.uk/ena/data/view/SRR493366) and for the fruit fly simulation we used those from the sample SRR1501444 (http://www.ebi.ac.uk/ena/data/view/SRR1501444). These samples both represent paired-end sequencing experiments with a read length of 101 bp. The transcriptome catalogs used as the basis for the expression estimation and simulation were Ensembl GRCh37.71 for human (using only the chromosomes in the primary genome assembly) and BDGP5.70 for the fruit fly simulation. For both organisms, we restricted the simulation to protein coding genes. Applying the RSEM expression estimation to the real data files provided us with a model file (detailing the sequencing settings and error model) and an isoform summary file (containing, among other things, the estimated TPM, expected count and relative abundance for each isoform). We modified the model file slightly to avoid simulating reads where most bases are of very low base quality.

### TPM estimation

We used the isoform-level count and TPM estimates provided by RSEM as the basis for the generation of sample-wise TPM values for the six samples. First, we summed the estimated counts for all isoforms of each gene, scaled to the desired library size (40 million for human and 25 million for fruit fly) and used a mean-dispersion relationship derived from real data (see [2]) to generate a matching negative binomial (NB) dispersion value for each gene. The gene count for each sample was sampled from a NB distribution with the estimated mean and dispersion parameters. We also calculated the gene-level read count per kilobase (RPK) by dividing the simulated read count by the effective gene length estimated by RSEM. Next, we simulated relative isoform abundances for each sample using a Dirichlet distribution with parameters set to the isoform fractions estimated by RSEM multiplied by a scale factor of 100. We selected 1,000 genes to be affected by differential isoform usage between the two conditions. These genes were selected randomly among the genes with at least two expressed isoforms (with relative isoform abundance above 10%) and a high enough expression level (expected gene count above 500). For each of these 1,000 genes, the relative isoform abundances of the two most abundant isoforms were reversed in the second condition (samples 4-6). This type of “switch events” was studied in detail in a previous publication [32] where it was shown that many genes underwent this type of modification between conditions. From the isoform percentages and the previously estimated gene RPKs, we estimated the isoform RPKs by multiplication. Finally, the isoform TPM was obtained by scaling of the RPK value. These transcript TPM values were used as the input to RSEM for simulating fastq files.

### Generation of fastq files and mapping

Given the TPM estimates for the individual samples and the modified RSEM model file, we used the `rsem-simulate-reads` module of RSEM to generate paired fastq files for the six samples. We simulated 40 million read pairs for each human sample and 25 million pairs for each of the fruit fly samples. Of these reads, 5% were simulated to come from a non-specific “background” (and thus not stem from any transcript). The fastq files were aligned to the reference genome and transcriptome using TopHat (v2.0.14) [33], provided with the reference gtf file.

### Incomplete annotation

To mimic the situation where the full transcriptome catalog of a studied organism is not known, we generated incomplete reference gtf files by excluding 20% of the transcripts (proportionally distributed between differentially used and non-differentially used). The entire data analysis pipeline, starting from the TopHat alignment, was re-run with these incomplete gtf files (using the same set of simulated data files).

### Detection of differential counting bin usage

Each of the counting methods applied in this study (see Additional file 1 for a more detailed description) generates a count matrix, where each row corresponds to a counting bin and each column corresponds to one of the six samples. They also contain information grouping the counting bins together based on their gene of origin. Each count matrix is submitted to the DEXSeq R package (v1.14.0) to test for changes in *relative usage* of each counting bin between the two simulated conditions. A difference in relative counting bin usage is interpreted as a preferential inclusion or exclusion of that bin in one condition compared to the other, which we interpret as evidence in favor of differential isoform usage between the conditions. More technically, for each counting bin, given the number of reads assigned to the bin and the sum of the reads assigned to all other bins of the gene in each of the samples, DEXSeq fits generalized linear models to test for the presence of an interaction between the condition and the “counting bin” factor (this bin *vs* all others). The bin counts are assumed to follow a negative binomial distribution and the dispersion parameters are estimated from the data and subjected to shrinkage using the method implemented in the DESeq2 R package [34]. Finally, a p-value for each bin summarizes the evidence in favor of differential usage of the bin between the conditions.

Given the statistical test results for each counting bin, we use the `perGeneQValue` function from the DEXSeq package to summarize the results on the gene level. This function associates a q-value to each gene by examining the number of genes for which at least one bin null hypothesis is rejected. DEXSeq applies an independent filtering step (adapted from DESeq2) and excludes counting bins with too low expression values from the testing. Genes for which all bins are filtered out are not assigned a q-value and are therefore not included in our evaluations. Depending on the number and structure of the counting bins, the collections of genes for which DEXSeq can assign a q-value differ between the counting methods. However, this does not affect the calculation of the observed TPR and FDR, since the excluded genes are not called differential.

Similarly to DEXSeq, *rMATS* provides one p-value per evaluated event (there may be multiple events for the same gene). These sub-gene structure p-values were summarized into gene-level q-values, quantifying the statistical evidence in favor of any differential usage of the counting bins of the gene, using the `perGeneQValue` function from DEXSeq. For *cuffdiff,* we used the gene-wise FDR estimates from the `cds.diff` output file. Importantly, this file records only differences in coding output between conditions and genes where the coding sequences of the differentially used isoforms are identical will therefore not be detected with *cuffdiff*.

## Availability of supporting data

The data sets supporting the results of this article are available in the ArrayExpress repository with accession number E-MTAB-3766 [http://www.ebi.ac.uk/arrayexpress/experiments/E-MTAB-3766/]. All code needed to reproduce the results are available from the project GitHub repository: https://github.com/markrobinsonuzh/diff_splice_paper. The list of truly differential genes, as well as the final results (gene q-values) for each method, are available from the accompanying website: http://imlspenticton.uzh.ch/robinson_lab/splicing_comparison/. The webpage also holds a link to a Shiny app that can take these summary files and reproduce the main figures of the manuscript.

## Competing interests

The authors declare that they have no competing interests.

## Authors’ contributions

CS and KLM designed analyses, analyzed data and drafted the manuscript. MN and CWL evaluated various pipelines and analyzed data. MDR designed the overall study and drafted the manuscript. All authors read and approved the final manuscript.

## Acknowledgements

The authors wish to thank Alejandro Reyes and members of the Robinson lab for helpful discussions and careful reading of the manuscript. We wish to acknowledge funding from: an SNSF Project Grant (143883), the European Commission through the 7th Framework Collaborative Project RADIANT (Grant Agreement Number: 305626), the SNSF-funded NCCR in RNA and Disease.

## Supplementary Materials

### Additional file 1 — Supplementary Figures, Tables and Methods (.pdf)

Additional file 1 contains Supplementary Figures and Tables referred to in the text. It also contains a description of the counting methods and a comparison of the isoform-guided filtering with bin-level filtering.

